# Exploring contributors to variability in estimates of SNP-heritability and genetic correlations from the iPSYCH case-cohort and published meta-studies of major psychiatric disorders

**DOI:** 10.1101/487116

**Authors:** Andrew J. Schork, David M. Hougaard, Merete Nordentoft, Ole Mors, Anders D. Børglum, Preben Bo Mortensen, Naomi R. Wray, Thomas Werge, for the iPSYCH Consortium

## Abstract

As more and more large psychiatric genetic cohorts are becoming available, more and more independent investigations into the underlying genetic architecture are performed, and an expanding set of replicates for estimates of key genetic parameters, namely, liability scale SNP heritability and genetic correlations – is amassing in the literature. In recent work, we published a set of SNP-heritability and genetic correlation estimates for major psychiatric disorders using data from the iPSYCH case-cohort study, and presented them alongside estimates gleaned from large, independently collected, analyzed and published meta-studies of the same disorders. Although in the broadest sense the estimates from iPSYCH and external meta-studies were concordant, and requiring strict statistical significance could reject the null hypothesis for few pairs, there were enough subtle trends in the differences to warrant further investigation. In this work, we consider a set of factors related to sample ascertainment, including the lifetime risks for disorders for the sampled populations, the use of age censored or partially screened controls, the sampling of extreme cases and controls, and diagnostic error rates, and attempt to assess their potential contributions to estimates of genetic parameters in the context of the difference trends observed in our previous work.

## Introduction

The underlying theory for the estimation of heritability on the liability scale^1-3^ is based on a model that defines the case-control status of individuals in the study as simple partitioning of a sample from a single population into two groups (cases and controls) at a threshold in unobserved liability that is defined by the lifetime risk for becoming a case. Whether real disease data conform to this assumption cannot be proven, but with data currently available the model seems robust and cannot be rejected. Given the model, transformations can be made from statistics estimated from the data to generate estimates of parameters on the liability scale, which at least allow benchmarking under a comparable scale across different scenarios. However, sometimes data collected for GWAS may be ascertained in such a way that they violate the assumptions inherent in the transformations. A series of recent papers have shown these deviations can introduce bias under the liability-scale SNP heritability framework, if the ‘standard’ transformations are applied, and provide revised transformations to account for sample ascertainment. In the following we discuss the implications of these papers to provide a qualitative investigation into the bounds on potential contributors to differences trends observed in the genetic parameters presented in Schork et al^4^.

### Baseline Difference Trends in Published Genetic Parameters

In Tables 1 and 2 we briefly summarize describe the key estimates presented in Figure 1 of Schork et al^4^ which we examine under a closer lens throughout, and their underlying sample sizes. All estimates in these tables have been published previously in Schork et al^4^, which itself culled GCTA GREML estimates of SNP heritability for external meta-studies from Lee et al^5^. LD-Score regression (LDSC)^6,7^ estimates from Schork et al^4^ used published meta-study GWAS summary statistics that have been studied widely: eXDX^8^, eADHD^9^, eAFF^10^, eANO^11^, eASD^12^, eBIP^13^, and eSCZ^14^. We present these published estimates to simply note the presence of some difference trends in estimates of genetic parameters in iPSYCH and published meta-studies for the same disorder or disorder pairs. What follows is an assessment of the potential contribution of ascertainment related factors to these observed (potential) differences.

**Table 1.**
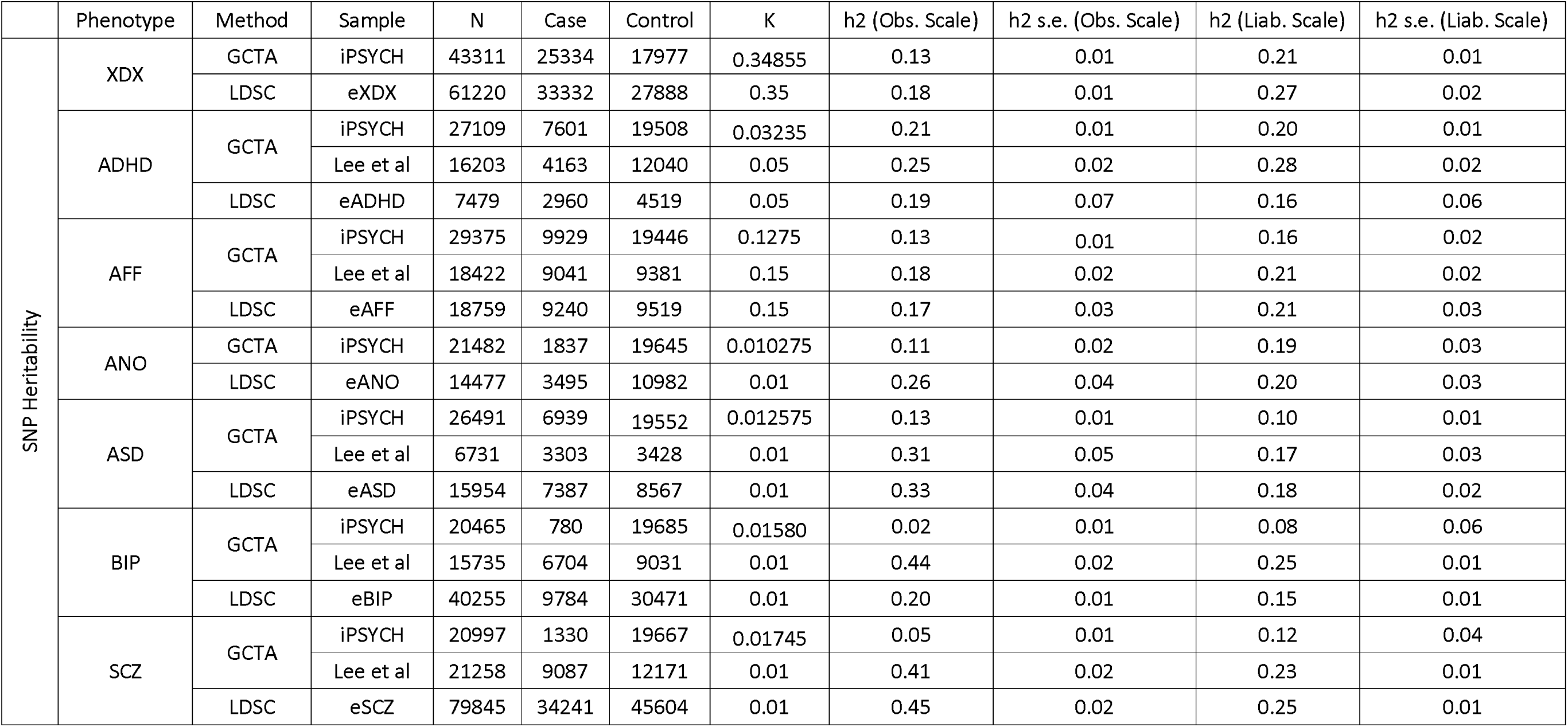
Estimates of SNP-heritability described in Figure 1 of Schork et al^4^. XDX, cross-disorder; ADHD, attention-deficit hyperactivity disorder; AFF, affective disorder; ANO, anorexia; ASD, autism-spectrum disorder; BIP, bipolar disorder; SCZ, schizophrenia; LDSC, LD-score regression; GCTA, GREML approach implemented in the GCTA software package; K, lifetime prevalence used for liability scale transformations; Obs., observed; Liab., liability; h2, SNP-heritability; s.e., standard error.

**Table 2.** Estimates of SNP-genetic correlation described in Figure 1 of Schork et al^4^. XDX, cross-disorder; ADHD, attention-deficit hyperactivity disorder; AFF, affective disorder; ANO, anorexia; ASD, autism-spectrum disorder; BIP, bipolar disorder; SCZ, schizophrenia; LDSC, LD-score regression; GCTA, GREML approach implemented in the GCTA software package; rG, SNP-genetic correlation; s.e., standard error.

|  | Phenotype 1 | Phenotype 2 | Method | Sample<br>(Phenotype 1) | Cases<br>(Phenotype 1) | Controls<br>(Phenotype 1) | Sample<br>(Phenotype 2) | Cases<br>(Phenotype 2) | Controls<br>(Phenotype 2) | rG | rG s.e. |
| --- | --- | --- | --- | --- | --- | --- | --- | --- | --- | --- | --- |
| SNP Genetic Correlation | ADHD | AFF | GCTA | iPSYCH | 6895 | 9635 | iPSYCH | 9223 | 9634 | 0.49 | 0.08 |
|  |  |  | LDSC | eADHD | 2960 | 4519 | eADHD | 9240 | 9519 | 0.43 | 0.18 |
|  | ADHD | ANO | GCTA | iPSYCH | 7538 | 9724 | iPSYCH | 1774 | 9725 | 0.01 | 0.09 |
|  |  |  | LDSC | eADHD | 2960 | 4519 | eADHD | 3495 | 10982 | 0.10 | 0.15 |
|  | ADHD | ASD | GCTA | iPSYCH | 6137 | 9693 | iPSYCH | 5474 | 9694 | 0.29 | 0.08 |
|  |  |  | LDSC | eADHD | 2960 | 4519 | eADHD | 7387 | 8567 | -0.09 | 0.14 |
|  | ADHD | BIP | GCTA | iPSYCH | 7485 | 9745 | iPSYCH | 664 | 9744 | 0.34 | 0.21 |
|  |  |  | LDSC | eADHD | 2960 | 4519 | eADHD | 9784 | 30471 | 0.39 | 0.12 |
|  | ADHD | SCZ | GCTA | iPSYCH | 7459 | 9737 | iPSYCH | 1188 | 9736 | 0.59 | 0.17 |
|  |  |  | LDSC | eADHD | 2960 | 4519 | eADHD | 34241 | 45604 | 0.23 | 0.08 |
|  | AFF | ANO | GCTA | iPSYCH | 9406 | 9700 | iPSYCH | 1314 | 9701 | 0.43 | 0.12 |
|  |  |  | LDSC | eAFF | 9240 | 9519 | eAFF | 3495 | 10982 | 0.35 | 0.11 |
|  | AFF | ASD | GCTA | iPSYCH | 9315 | 9654 | iPSYCH | 6325 | 9653 | 0.61 | 0.11 |
|  |  |  | LDSC | eAFF | 9240 | 9519 | eAFF | 7387 | 8567 | 0.24 | 0.12 |
|  | AFF | BIP | GCTA | iPSYCH | 9929 | 9713 | iPSYCH | 780 | 9713 | 0.82 | 0.34 |
|  |  |  | LDSC | eAFF | 9240 | 9519 | eAFF | 9784 | 30471 | 0.48 | 0.08 |
|  | AFF | SCZ | GCTA | iPSYCH | 9462 | 9709 | iPSYCH | 863 | 9709 | 0.96 | 0.28 |
|  |  |  | LDSC | eAFF | 9240 | 9519 | eAFF | 34241 | 45604 | 0.51 | 0.05 |
|  | ANO | ASD | GCTA | iPSYCH | 1749 | 9747 | iPSYCH | 6851 | 9747 | 0.29 | 0.12 |
|  |  |  | LDSC | eANO | 3495 | 10982 | eANO | 7387 | 8567 | 0.03 | 0.09 |
|  | ANO | BIP | GCTA | iPSYCH | 1810 | 9813 | iPSYCH | 753 | 9812 | 0.15 | 0.23 |
|  |  |  | LDSC | eANO | 3495 | 10982 | eANO | 9784 | 30471 | 0.29 | 0.06 |
|  | ANO | SCZ | GCTA | iPSYCH | 1791 | 9805 | iPSYCH | 1284 | 9804 | 0.47 | 0.21 |
|  |  |  | LDSC | eANO | 3495 | 10982 | eANO | 34241 | 45604 | 0.25 | 0.04 |
|  | ASD | BIP | GCTA | iPSYCH | 6901 | 9766 | iPSYCH | 742 | 9766 | 0.67 | 0.33 |
|  |  |  | LDSC | eASD | 7387 | 8567 | eASD | 9784 | 30471 | 0.15 | 0.08 |
|  | ASD | SCZ | GCTA | iPSYCH | 6825 | 9758 | iPSYCH | 1216 | 9758 | 0.73 | 0.23 |
|  |  |  | LDSC | eASD | 7387 | 8567 | eASD | 34241 | 45604 | 0.21 | 0.06 |
|  | BIP | SCZ | GCTA | iPSYCH | 711 | 9824 | iPSYCH | 1261 | 9824 | 0.59 | 0.39 |
|  |  |  | LDSC | eBIP | 9784 | 30471 | eBIP | 34241 | 45604 | 0.75 | 0.03 |

**Figure Set 1.**
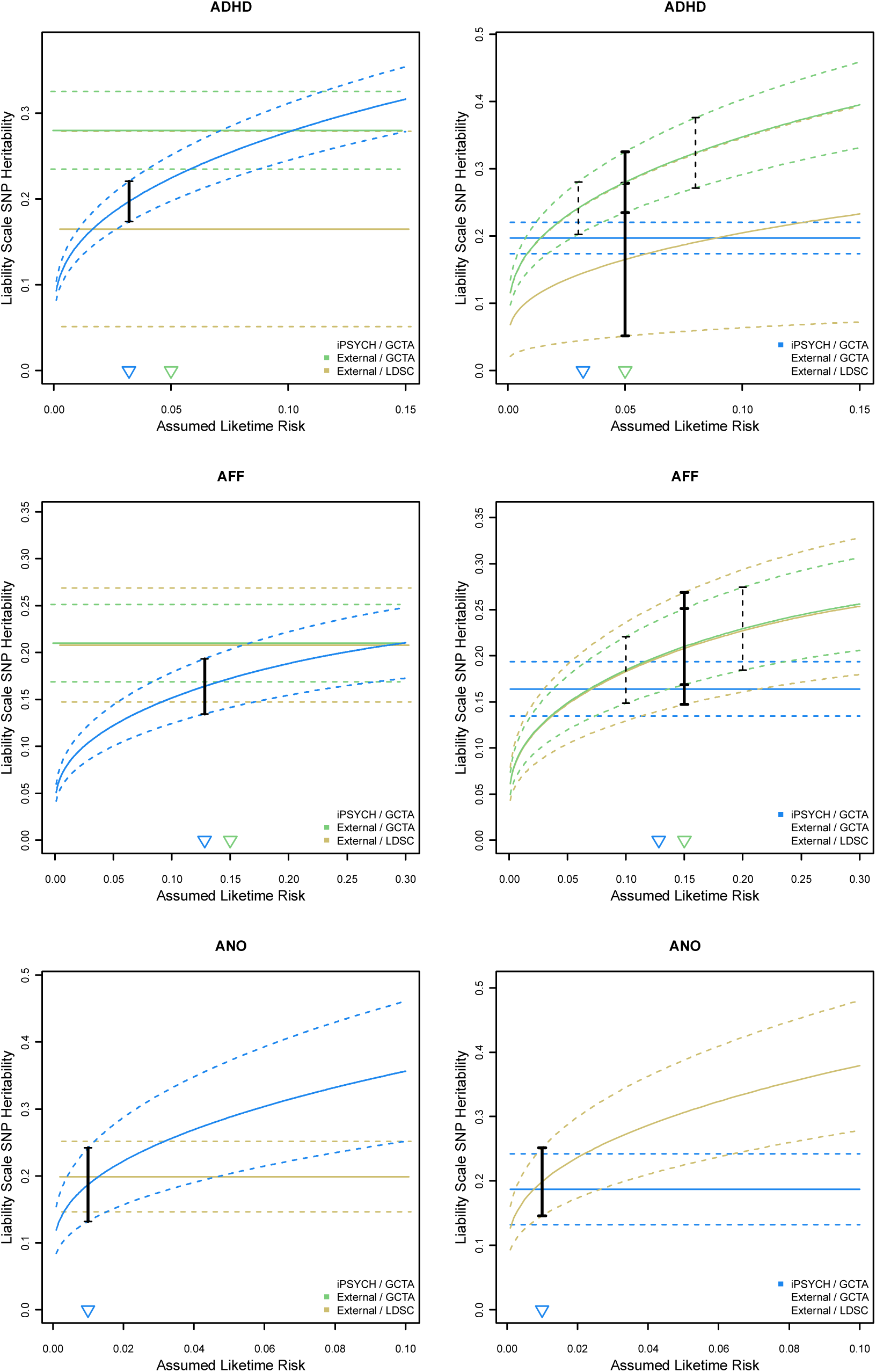
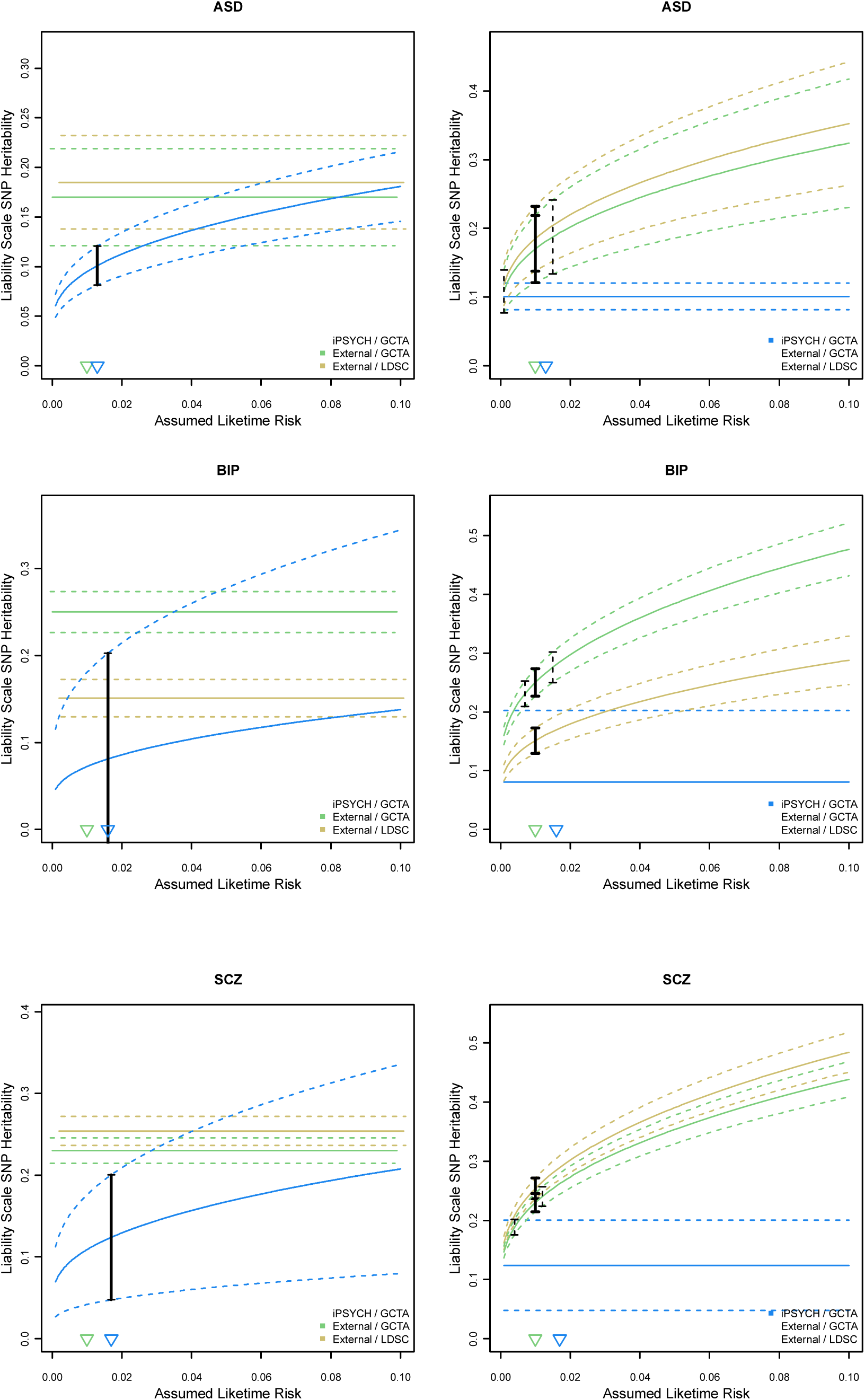
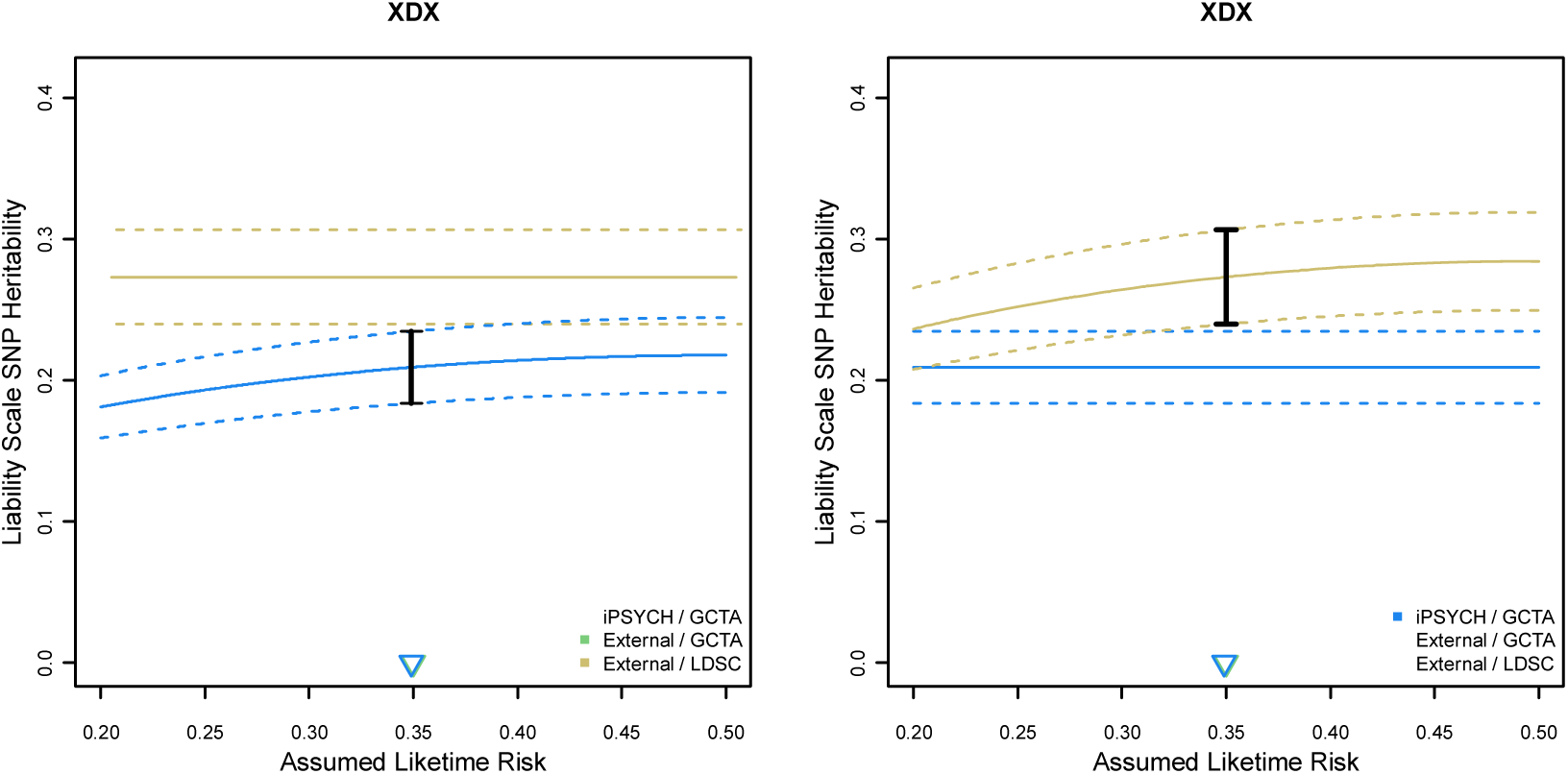
Effects of different assumed lifetime risk on estimates of liability scale heritability. Each pair of figures (a row) depicts the estimates of SNP-heritability from Schork et al^4^ described in table 1. The figure on left shows the iPSYCH estimate of SNP-heritability re-computed for a range of assumed lifetime risk (solid blue curve) and it’s approximate 95% confidence interval (dashed blue curve). The solid black bar shows the estimate and 95% confidence interval when assuming the lifetime risk estimated in the Danish population by Pedersen et al^17^. The pale yellow and pale green rectangles show the 95% confidence regions for the external estimates taken from GCTA (Lee et al^15^) and computed for external GWAS using LDSC regression, respectively, at the prevalence described in Table 1. In the right figures for each disorder, pale yellow and pale green curves show the GCTA and LDSC regression external estimates and 95% confidence intervals over a range of assumed lifetime risk, while the blue rectangle shows the 95% confidence interval for the iPSYCH estimate at the assumed lifetime risk described in Table 1. Solid dots and vertical bars represent the external estimates at the assumed risk from Table 1. For data taken from Lee et al, we also include dashed vertical bars to denote their specified plausible bounds of lifetime risk in the previous study. Blue and green triangles along the x-axis mark the lifetime risk from table 1 for iPSYCH and external estimates, respectively.

### Estimates of Population Lifetime Risk

Whether estimated by GREML or LD-score regression, SNP-heritability^15^ is the proportion of phenotypic variance explained by common SNP-variation from unrelated subjects. For quantitative traits the phenotypic variance is estimated well from a sample drawn from the population. For binary case-control (0/1) data the phenotypic variance in a sample is P(1-P), where P is the proportion of the sample that are cases, or the lifetime risk. Hence, the estimate for the variance explained by all SNPs is made on the “observed scale” (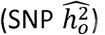). This estimate is then transformed to be an estimate on the liability scale (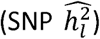) according to parameters describing the underlying population distribution in liability: 1) K, the lifetime risk of the disorder and 2) z, the height of the standard normal curve at the liability quantile (the case threshold) corresponding to a tail probability of K, and here assuming that our case-control sample has been randomly drawn from the population and everyone in the population who will get the disease has the disease, P=K (i.e. no age censoring).

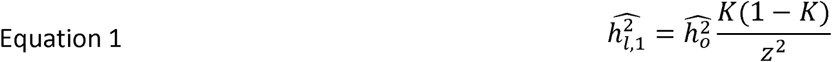

Often GWAS data are not consistent with the assumptions of this simple transformation because they are typically ascertained for cases at levels well above the proportion expected given the population lifetime risk, which will produce biased (i.e., inflated or deflated) estimates^15,16^. Lee et al^15^ propose corrections to the transformation applied to estimates of 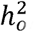, the magnitude of which depends on P for the sample. To emphasize that the estimated 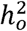 is dependent on the properties of the case-control sample, rather than the population, the estimate from ascertained data is called 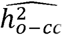. The updated transformation accounts for ascertainment through its effect on the variance in the population K(1-K) relative to the variance in the sample P(1-P).

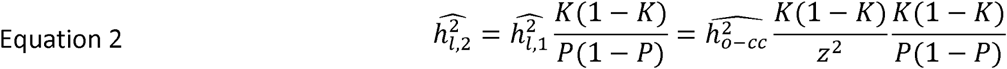

If an inaccurate lifetime risk is used for the above transformation, then an inaccurate heritability estimate will result. True lifetime risk estimates for a sampled population can be surprisingly hard to find, and so sensitivity analyses can be conducted considering a plausible range of K. The impact is usually trivial when K is 0.01 (with a plausible range of 0.005 to 0.02) relative to the assumptions of the model, but can become less trivial when K is 0.1 with a plausible range from 0.05 to 0.15. Although in Lee et al^5^, the study from meta-study GCTA estimates were obtained, this is discussed, errors in estimates of K may not be regularly accounted for a source of variability in estimates, intuitively or statistically (e.g. via adjusted standard errors). Estimating heritability in meta-studies aggregating multiple case-control samples from different populations using different case definitions requires assuming a risk that is not well defined, limiting the objective precision of SNP-heritability estimates. iPSYCH has the unique advantage of published lifetime risk estimates^17^ calculated for the same diagnostic criteria and within the same population SNP-heritability is studied in (XDX, 0.349; ADHD, 0.032; AFF, 0.128; ANO, 0.010; ASD, 0.013; BIP, 0.016; SCZ, 0.017), which provide similar, but not identical numbers to those emphasized in Lee et al (ADHD, 0.05; AFF, 0.15; ASD, 0.01; BIP, 0.01; SCZ, 0.01). In Figure Set 1 below, we show the effect of using different lifetime prevalence estimates on the estimates of SNP-heritability for each diagnosis in iPSYCH and for the external meta-studies. These data show that, especially for rare diseases where the immediate slope of the liability heritability versus assumed lifetime risk curve is steepest, assuming the wrong lifetime risk just slightly could change the estimate of heritability by as much as a standard deviation (half of the confidence interval). The iPSYCH data is linked with unambiguously appropriate estimates of lifetime risk, although these are still subject to sampling variance^3^ of K(1-K)/z^2^N, where N is the size of the population from which the estimates are made. So, for a single birth cohort year from Denmark (population 5.7M, ~61,000 births per year), the standard error on the estimate of K =0.01 is 0.015, but reducing to a standard error of 0.0015 when 10 years of birth cohort data are used. For the external meta-studies, accurate estimates of K are not available, as the aggregated data may come from different populations, with different phenotype definitions, and thus different lifetime risks, introducing uncertainty. In Lee et al^5^, the plausible ranges of K they consider can result in estimates of heritability for the upper and lower bounds of K that are as much as two standard deviations apart (the width of the confidence intervals barely overlap). This additional source of variability, which is not typically reflected in the estimated standard errors and confidence intervals, and the likelihood of its effect in both cohorts should be considered intuitively when thinking about estimates of heritability from iPSYCH and external studies. Estimation of genetic correlations are scale independent^15^, so these estimates are not likely to be susceptible to misestimation of K.

### Use of (Partially) Screened Controls

One assumption of the standard transformation equation of SNP-heritability estimates from case-control to liability scale, i.e., 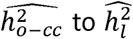 is that control samples are completely screened for the condition being studied^18^. This is not always the case, for example when a sample from the general population is used as controls and diagnoses are unavailable. The work by Peyrot et al^18^ shows how using unscreened controls leads to a downward bias in the estimation of heritability and derives a correction factor. The correction adjusts for the proportion of the controls that are expected to be unscreened cases, which they call false controls (F = N_false controls_ / (N_false controls_ + N_true controls_)) and extends the transformation above.

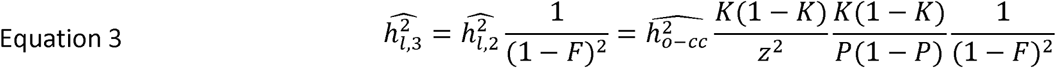

In addition, if control subjects are young enough that they have not expressed their lifetime risk, then they can be viewed as “partially screened” – they have been screen for disease *only up to their current age*. In a large enough sample of young individuals, a (substantial) proportion will be expected to develop a disorder later in life. This could be especially pertinent for the iPSYCH cohort given its youth (all controls born between 1981 and 2005) relative to external studies which may employ older controls.

We can again take advantage of the extensive epidemiological work that preceded the iPSYCH study^17^, which not only provides estimates lifetime risk, but also gender specific cumulative risk estimates across the lifespan (Supplementary Tables of Pedersen et al^17^). We leverage these estimates to explore the potential under-estimation of heritability in iPSYCH due to the youth of the subjects, assuming the framework proposed in Peyrot et al^18^. For each control subject, we define their remaining risk (K_age+_) for a disorder as the lifetime risk of the disorder (K), minus the cumulative risk up to their age (K_age-_), conditional on their gender. This number can be seen as a rough estimate of the probability that a given control will convert to being a case at some point in their life. For the control group for each disorder, we estimate the expected number of false controls (N_false controls_) by summing these probabilities across all control individuals for the disorder. We set a plausible range for these numbers by estimating the expected number of false controls using the upper and lower 95% confidence intervals for the lifetime cumulative risk estimates (Supplementary Tables of Pedersen et al^17^). We further extend this range to account the smaller sample in iPSYCH, relative to Pedersen et al. We extend the lower and upper bounds by two standard deviations of binomial distributions where the probability is defined the by the estimated number of false controls over the total number of controls (F) and the number of draws by the total number of controls. Our estimates for the expected number and proportion of false controls are described in Table 3 below.

**Table 3.** Estimates for the number and proportion of false controls in the iPSYCH data

|  | ADHD | AFF | ANO | ASD | BIP | SCZ | XDX |
| --- | --- | --- | --- | --- | --- | --- | --- |
| N<sub>False Cases</sub> | 206.66 | 2084.93 | 100.42 | 41.82 | 284.21 | 263.15 | 4414.25 |
| Range Lower Bound | 176.23 | 1988.09 | 79.42 | 28.3 | 246.62 | 228.06 | 4291.69 |
| Range Upper Bound | 236.13 | 2181.84 | 122.88 | 54.9 | 323.31 | 299.38 | 4536.67 |
| F | 0.0106 | 0.1072 | 0.0051 | 0.0021 | 0.0144 | 0.0134 | 0.2455 |
| F Lower Bound | 0.009 | 0.1021 | 0.004 | 0.0014 | 0.0125 | 0.0116 | 0.2386 |
| F Upper Bound | 0.0121 | 0.1123 | 0.0063 | 0.0028 | 0.0165 | 0.0153 | 0.2525 |

In Figure Set 2 below we describe the potential effects of having a relatively young control cohort on our estimates of SNP heritability. These data show that in general the youth of the iPSYCH cohort is not expected to have introduced substantial downward bias in our estimates of SNP-heritability, as even in the worst-case scenario, the prevalence of any single mental disorder is not sufficient to result in large numbers of false controls. An exception for this trend may be for AFF, where the prevalence is relatively higher and there is an appreciable proportion of risk that appears late in life^17^. The effect on XDX estimates also appear large, however, these numbers may be less accurate because an implicit assumption of the correction factor is that the misclassified cases are equivalent genetically to the identified cases, which may not hold in this case. For a definition of XDX that is “all mental disorders,” this will not hold because our definition of XDX is enriched for the iPSYCH ascertained diagnoses (i.e., it is not perfectly representative of the population of all psychiatric patients). Also, any changes in genetic architecture, (e.g. the magnitude of heritability or levels of genetic heterogeneity) that are a function of age of onset would result in differences between the observed cases and false controls and would be expected to alter these descriptions.

**Figure Set 2.**
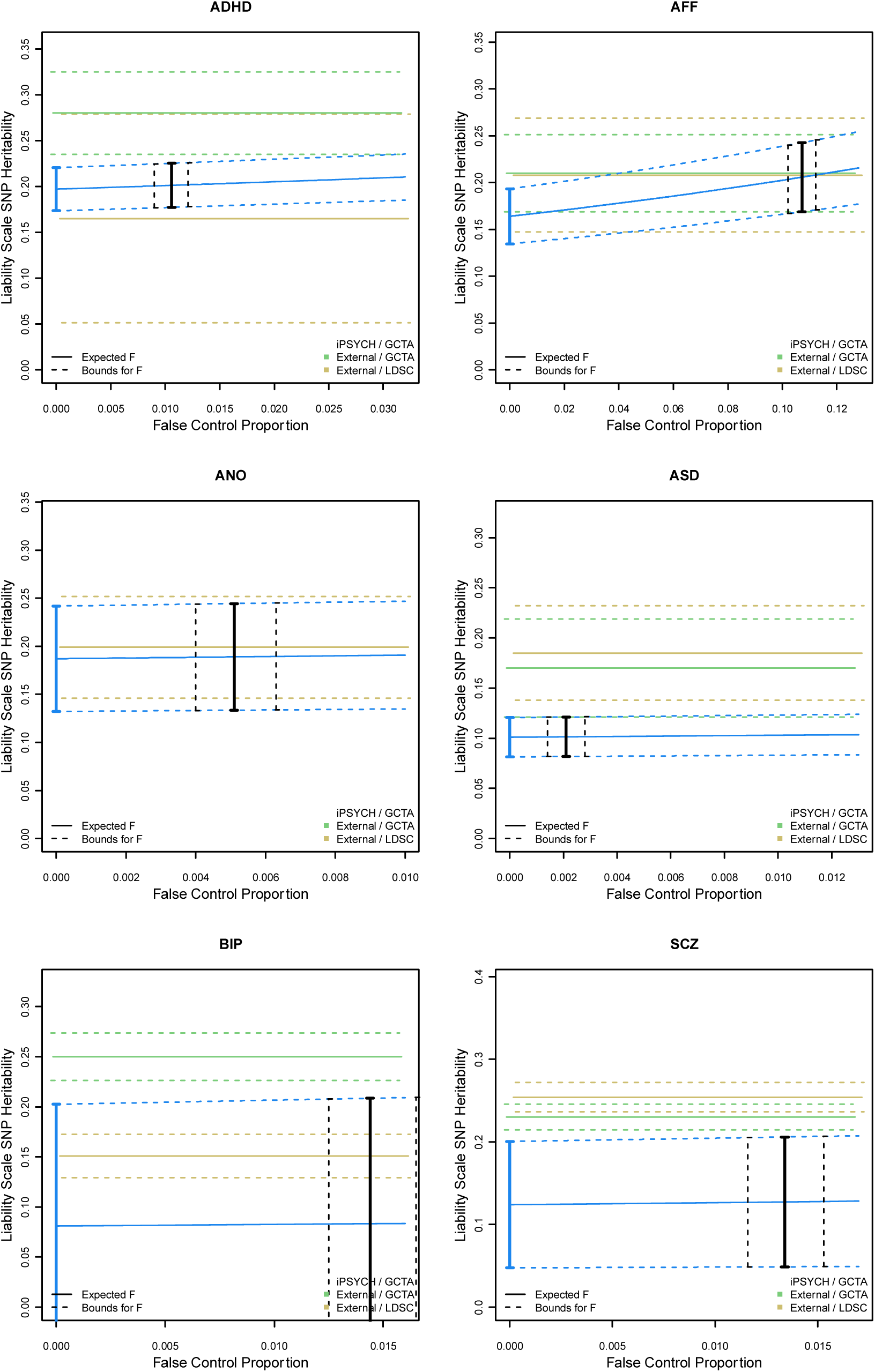
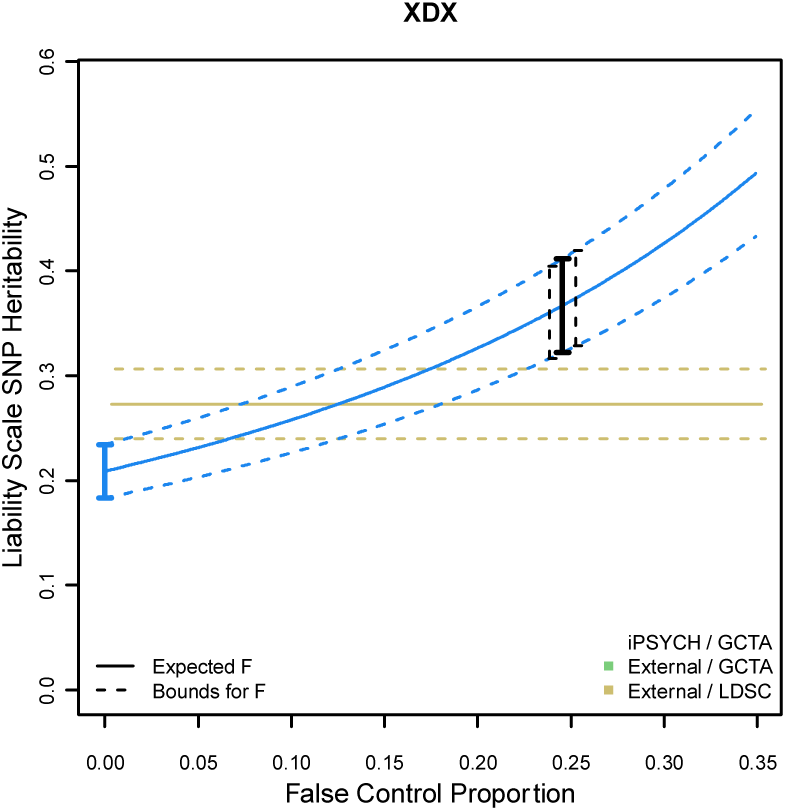
Bounding the effects of age censoring as misclassified controls. Each figure shows the estimated liability scale SNP-heritability (solid blue curve) and its 95% confidence interval (dashed blue curves), after accounting for a given proportion of false controls, ranging from 0 (fully screened) to the population life risk (fully unscreened). Blue dot and vertical bar represent the unadjusted estimate provided in Table 1, while the black solid bar and dashed bars provide the estimates given expected proportion of false controls and potential upper and lower bounds, respectively. Yellow and green rectangles show the 95% confidence intervals for the LDSC regression and GCTA external estimates of SNP-heritability, respectively, presented in Table 1.

Unfortunately, the necessary data for the external studies are not available to explore these transformations, but we would expect the effect to be reduced as iPSYCH is exceptional in its youth. This means the youth of iPSYCH would likely lead to exaggerated differences between iPSYCH and external studies of heritability (assuming the true values are the same). However, the contribution of this potential downward bias (which is already modest) is expected to be further tempered by the same (but less severe) age censoring in published data. We are not aware of a published framework for investigating the impact of unscreened or partially screened controls on estimates of SNP-based genetic correlation and see this as an avenue for future research.

### Extreme sampling of cases and/or controls

Another assumption of the standard transformation from 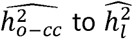 is that cases and controls are a *random sample* of a population’s cases and controls, with respect to the single liability partitioning. Data collected for GWAS, as used for heritability estimation in the external studies, may not only be ascertained for *extra* cases, but also often employ *extreme* cases and/or controls, which could violate this assumption. Extreme controls can be defined as ascertained control subjects with, on average, less genetic liability for the disorder being considered than would be expected from a random sample of the underlying population of unaffected subjects. In practice, this could arise when subjects are selected for an absence of family history, to be free of specific disorders that are genetically correlated with the outcome of interest, or for exceptional mental health, in general. It is known that in the Psychiatric Genetics Consortium (PGC, who published most of the external studies we cite), for example, some contributing cohorts use the same control individuals for multiple disorders or select for other aspects of psychiatric health within the control samples. Extreme cases can be similarly defined as ascertained cases with, on average, more genetic liability than would be expected in a representative sample of cases from the underlying population. In GWAS, cases may be ascertained for prevalent cases, treatment resistant cases, disease severity, or archetypical presentation. If any of these features are correlated with increased genetic liability, as, for example, has been recently suggested for schizophrenia^19,20^, an upward bias in genetic liability relative to a random selection cases is expected and it is expected to translate to an upward bias in heritability.

This issue was introduced formally in a recent commentary^21^ describing the mis-estimation of heritability for male pattern baldness that was a result of censoring cases of “rather dubious baldness.” Yap et al^21^ suggested that this censoring resulted in case and control populations that were extreme with respect to genetic liability by unintentionally censoring an intermediate portion of the underlying genetic liability distribution. As a result, the original paper^22^ produced a SNP-heritability estimate biased substantially upwards, by approximately 50% of the corrected value. To counter this, Yap et al (Supplementary Methods)^21^ derive a transformation for observed scale heritability that takes into account ascertainment of extreme cases and/or extreme controls by decoupling the lifetime risk of being a case from that of remaining a control. In the original transformations proposed by Lee et al^15^, cases represent the upper K proportion of the population liability and controls the lower 1-K proportion, reflecting the notion that the single partitioning of the underlying population is assumed to be mirrored in the study sample. Yap et al^21^, drawing on prior work of Gianola^23^ and Golan et al^16^, generalize this to reflect ascertainment of case and control samples that represent more extreme proportions of the underlying liability, which may include a “gap” due to intermediate individuals being excluded from the study. This model uses two independent thresholds on the underlying liability scale, defining the lifetime risk of being an extreme case (K_U_) and extreme control (K_L_) independently, with the definition of Z_U_ and Z_L_ the height of the standard normal at the quantiles defined by these tail probabilities. When K_u_ = K, and K_L_=1-K, their correction reduces to the equation from Lee et al^15^ above.

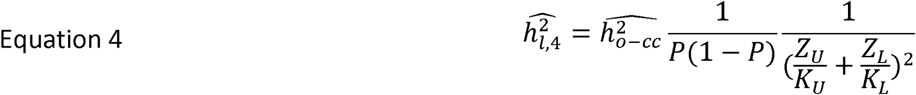

In iPSYCH, the data should be fairly, and perhaps uniquely, free from potential extreme sampling ascertainment bias because of how case and control populations have been drawn from the broader Danish population and aggregated for analysis (see Pedersen et al^24^ for a description of the sampling scheme). The iPSYCH groups were defined to mirror a simple, single partitioning of underlying population liability. Controls are random samples of the population of unaffected subjects, without additional censoring, and cases include all diagnosed individuals, regardless of features that may reflect genetic severity. The extent of extreme sampling bias in the external meta-studies is hard to determine because the contributing cohorts will have different criteria for inclusion, and the censoring of controls or ascertainment for cases may not be directly on the genetic liability for the disorder, as was the case for the baldness study, obscuring appropriate estimates for K_U_ and K_L_. To illustrate, consider screening schizophrenia controls additionally for bipolar disorder. Because of the high genetic correlation that is more or less accepted between these disorders, the remaining control population will be depleted for the portion of genetic risk that is shared between the disorders. In essence, this censors individuals that are expected to be intermediate with respect schizophrenia liability, similar in concept to the example from Yap et al. However, because the genetic correlation between the disorders is not one, they are not directly selected to be depleted of schizophrenia liability, and so the proportion of the population being represented by the screened controls is not simply 1-K_SCZ_-K_BIP_. The lower tail proportion (K_L_) of the population represented by sampling bipolar censored schizophrenia controls may be difficult to define but should certainly be less than the assumed 1-K_SCZ_. The same complications arise when oversampling severe, prevalent or archetypical cases. The proportion of the underlying population with respect to liability represented by ascertained subjects, K_U_, is surely less than K (i.e., they are likely more extreme with respect to underlying liability than an average case), but the correct choice for K_U_ requires knowing the relationship between the ascertained features and underlying liability. Investigating this fully in the context of psychiatric disorders requires additional data from external studies and the development of novel analytic paradigms. As such, we can only attempt to provide crude benchmarks for this effect in hope to motivate more thorough future work.

In order to qualitatively explore this phenomenon, we again take advantage of the ascertainment in iPSYCH, providing unselected cases and controls. We benchmark a plausible bound on the value for K_L_ in ascertained GWAS data, by estimating the SNP-heritability for each disorder after censoring the control population for all other psychiatric diagnoses, and comparing the resulting SNP-heritability with the estimate obtained when using the appropriate uncensored controls. In Figure Set 3 below (lightest grey bars) we show a universal overestimate, albeit modest, of heritability when using extreme controls created by censoring subjects with additional diagnoses, consistent with the concepts described in Yap et al^21^. By assuming the true heritability (h_I_^2^) is equal to the estimate computed with uncensored controls, and, due to the iPSYCH design where our cases should not be ascertained for higher than expected liability (i.e., K_U_=K), we can solve equation 4 above (taken from Yap et al^21^) for K_L_. This estimates the proportion of the population for, say, schizophrenia liability, that are being sampled from when controls are indirectly censored by excluding subjects with any other psychiatric diagnoses. We provide benchmark estimates of K_L_ for each disorder in Table 4, below. Consistent with intuition, the difference between K_L_ and 1-K when controls are censored for secondary conditions is largest for phenotypes with the largest genetic correlation with the remaining diagnoses (SCZ, BIP, AFF; see Schork et al^4^ Supplementary Figure 4) and smallest for those with less genetic correlation (ANO; see Schork et al^4^ Supplementary Figure 4).

**Figure Set 3.**
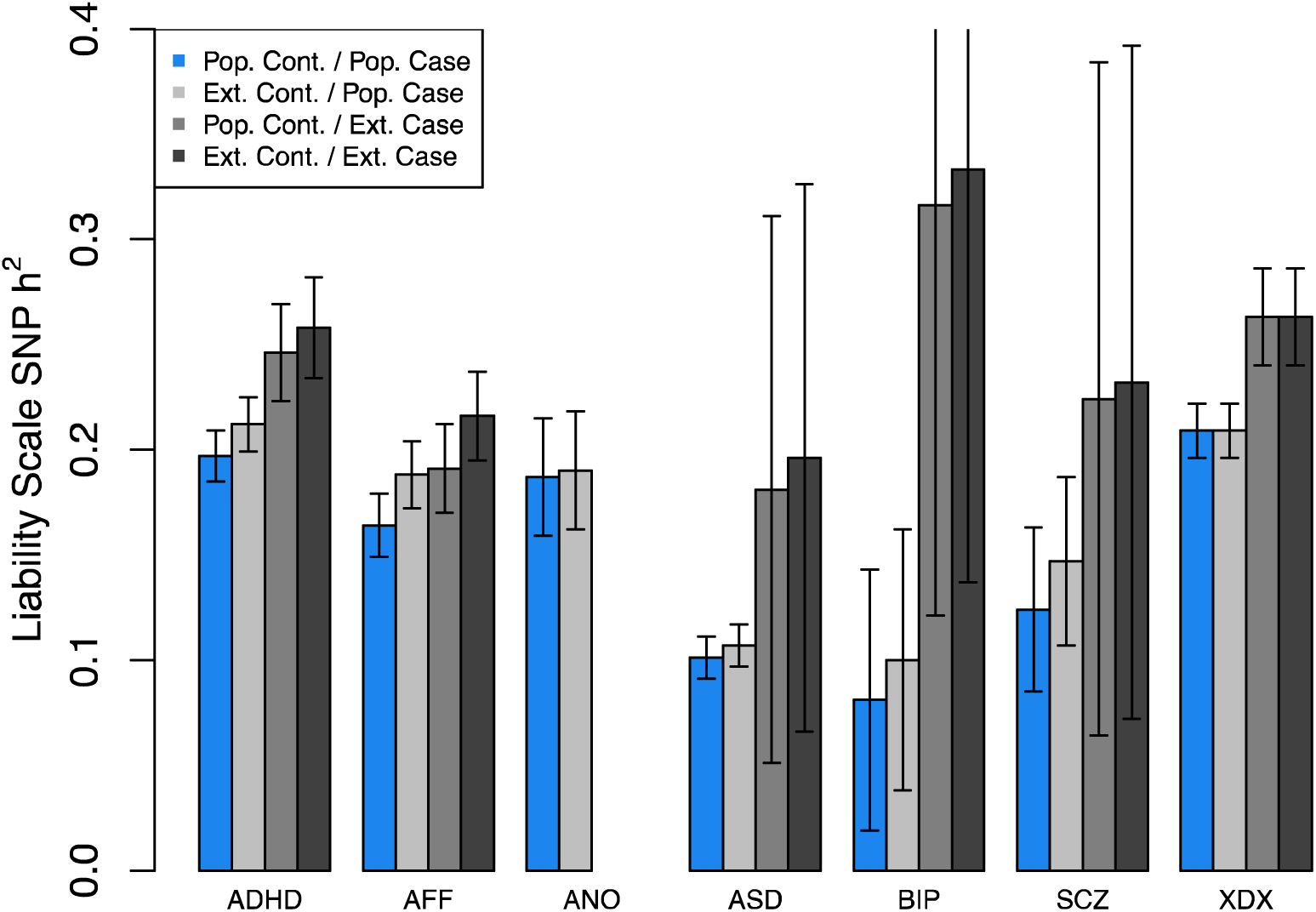
Re-estimation of iPSYCH SNP heritability after extreme sampling of cases and/or controls.

**Table 4.** Benchmark values K_L_ and K_U_ inferred indirectly from iPSYCH data. *the controls cannot be additionally censored for XDX, by definition of the phenotype, so this confirms that our back calculations return 1-K, modulo rounding error, work as intended. **Severity information for ANO was not currently available.

|  | ADHD | AFF | ANO | ASD | BIP | SCZ | XDX |
| --- | --- | --- | --- | --- | --- | --- | --- |
| Assumed 1-K | 0.968 | 0.872 | 0.99 | 0.987 | 0.984 | 0.983 | 0.651* |
| Plausible $K_L$<br>Lower Bound | 0.925 | 0.787 | 0.985 | 0.946 | 0.810 | 0.855 | 0.653* |
| $K_L / (1-K)$ | 0.956 | 0.903 | 0.995 | 0.958 | 0.822 | 0.870 | 1 |
| Assumed K | 0.032 | 0.128 | 0.010 | 0.013 | 0.016 | 0.017 | 0.349 |
| Plausible $K_U$<br>Upper Bound | 0.016 | 0.09 | NA** | 0.001 | 0.0001 | 0.001 | 0.256 |
| $K_U / K$ | 0.494 | 0.729 | NA** | 0.062 | 0.006 | 0.071 | 0.734 |

**Figure Set 4.**
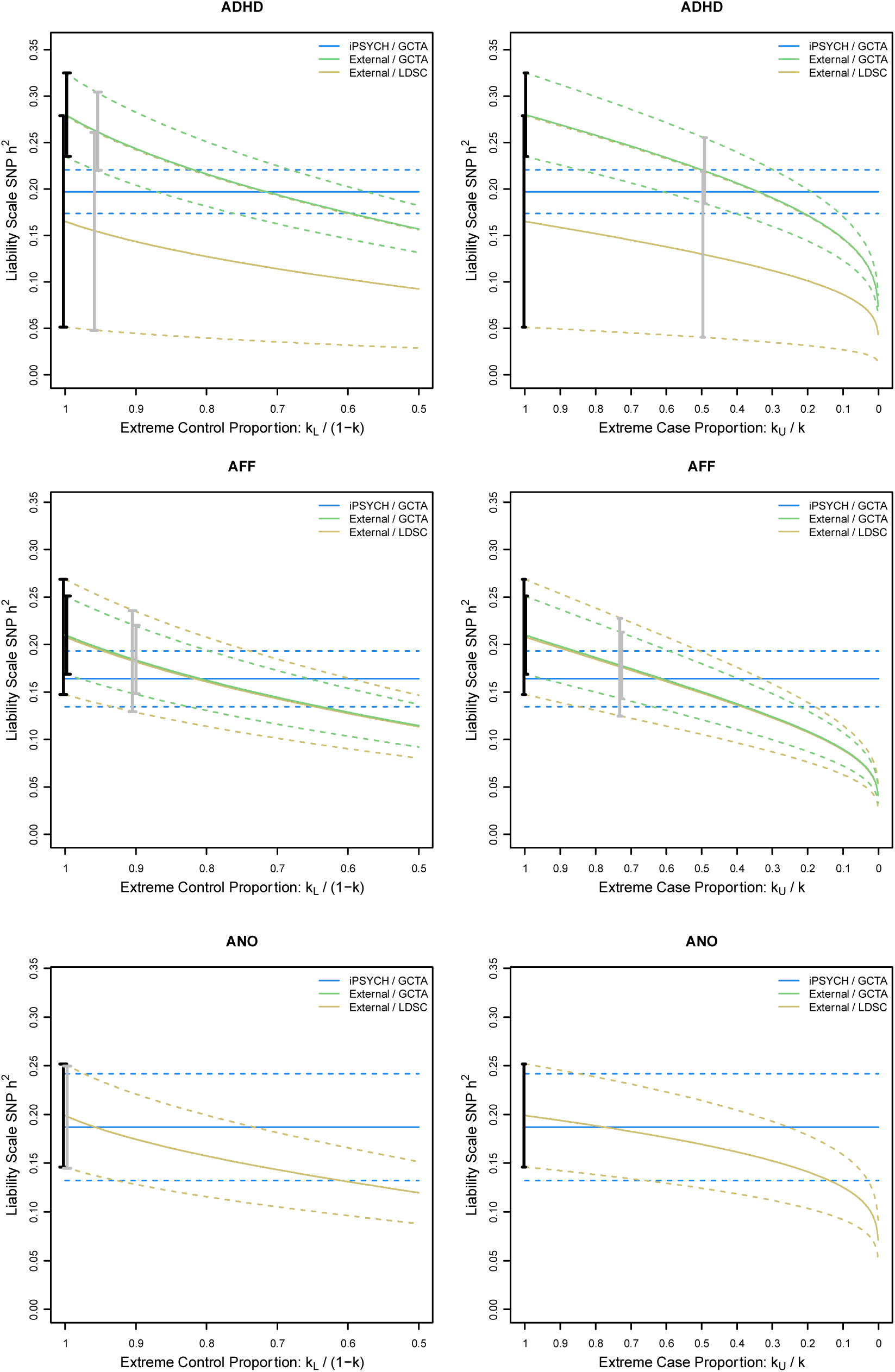
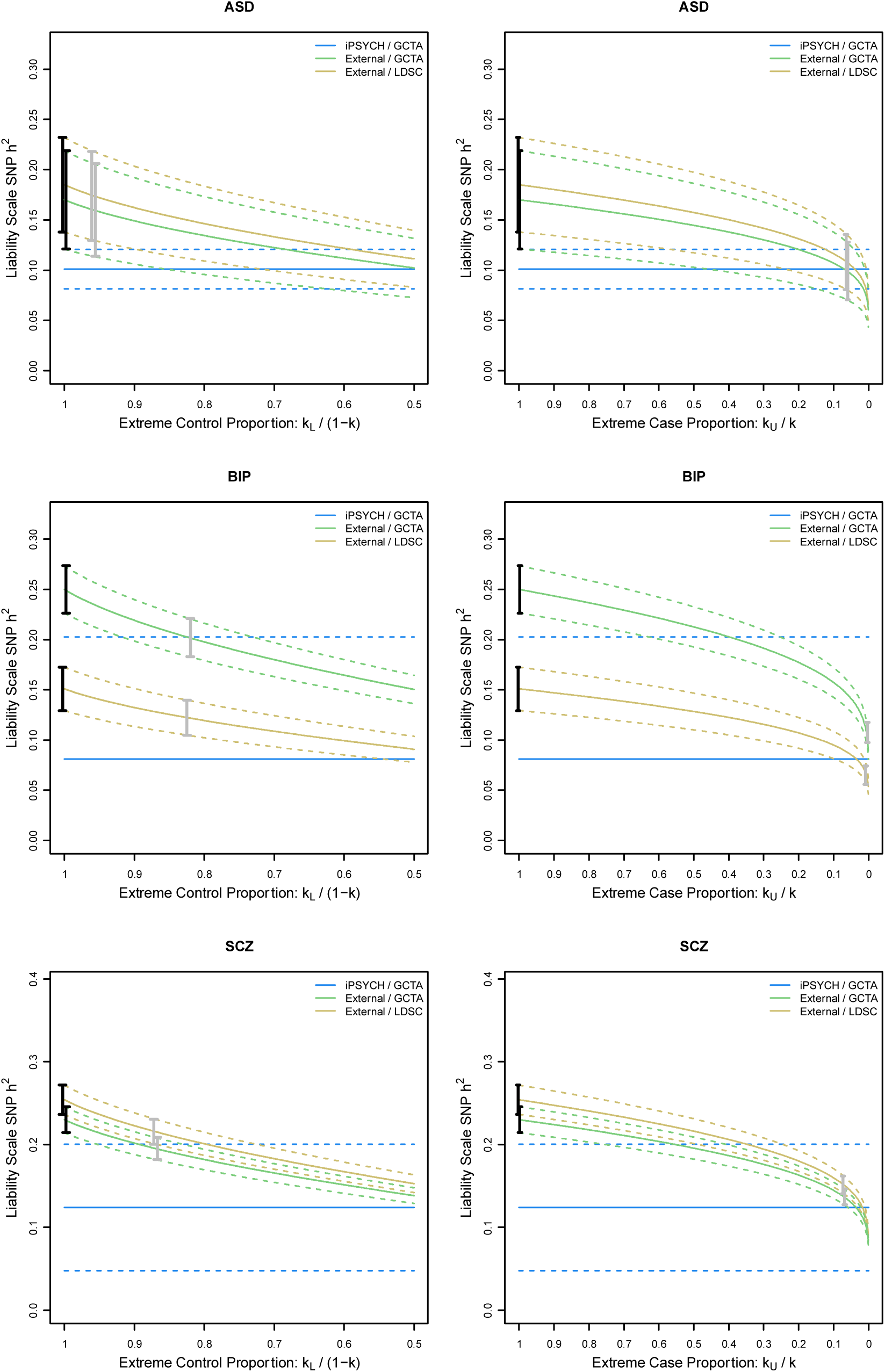
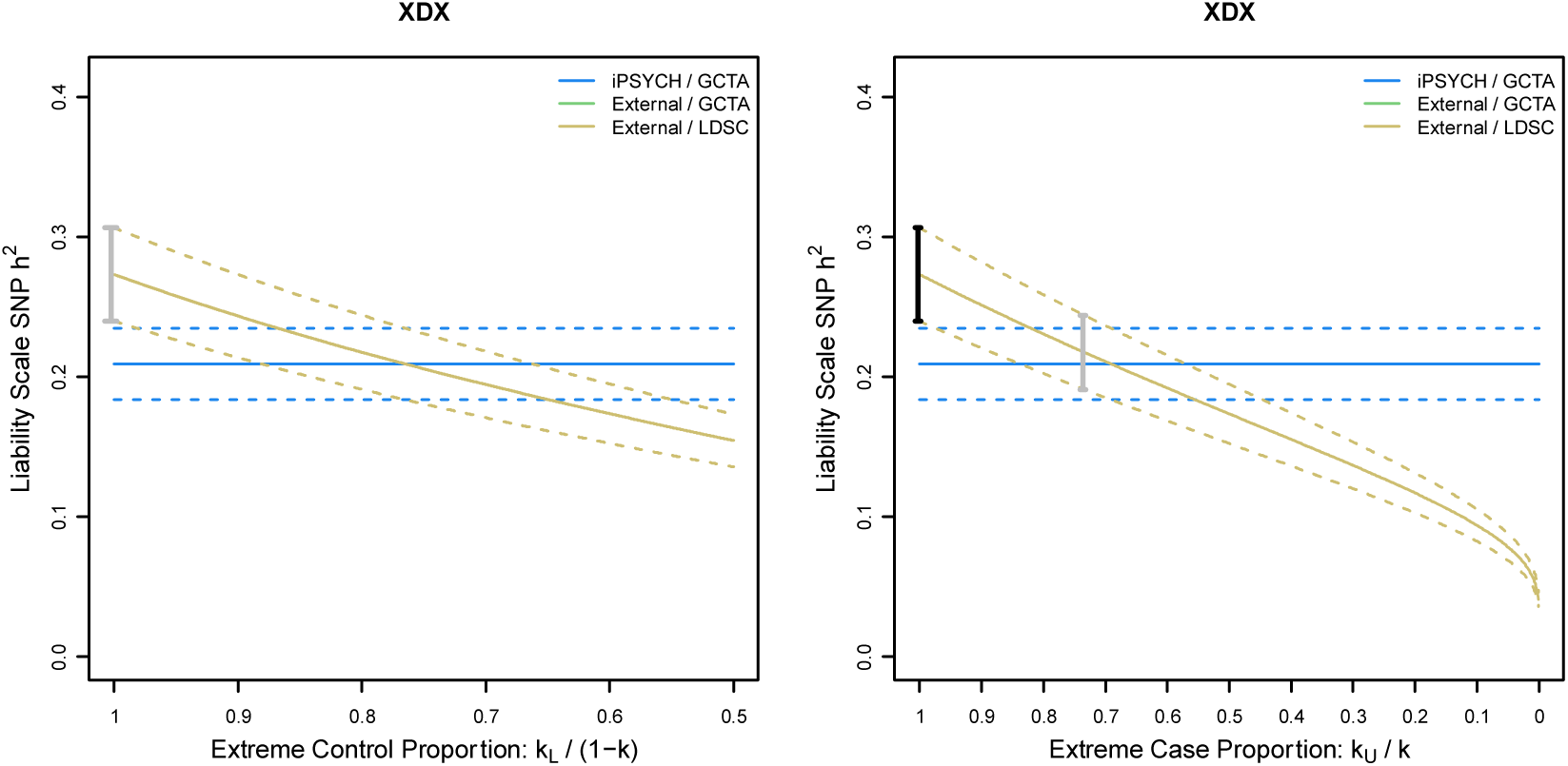
Exploring the effects of extreme sampling of either cases or controls. In each pair of figures (a row) we show the effects of correcting for extreme sampling on external estimates of SNP heritability for GCTA (green curves with 95% confidence intervals) and LDSC regression (yellow curves with 95% confidence intervals). The point estimate and 95% confidence intervals computed in iPSYCH are covered by the blue rectangle. The black dot and confidence interval shows the original estimate from Table 1, while the grey dot and confidence interval highlights the corrected value using K_L_ or K_U_ presented in Table 2, above.

To explore a possible bound on the values for K_U_ in ascertained GWAS data, we censored cases according to a plausible proxy for severity, the total number of hospital contacts an individual experienced for the disorder, and re-estimated SNP-heritability using the iPSYCH data. Frequency and length of hospitalization was recently shown to correlate positively with genetic liability for schizophrenia^19^ and it has been supposed that clinically ascertained research cohorts may be enriched for prevalent cases which are noted by more frequent and longer duration of hospital contacts, among other potentially exaggerating features^19,25^. Because the exact function defining the relationship between liability and this proxy for severity is unknown, and may vary across disorders, we estimated the SNP-heritability censoring cases for greater than the 0.25, 0.5 and 0.75 quantiles in number of hospital contacts (data not shown) and report the largest SNP-heritability from these three case definitions. This will surely over-fit the effect of censoring on hospital contacts, but provides us an estimate that we can use as an intuitive, albeit imperfect, attempt to provide a worst-case upper bound of the effect. As such, our estimates should be treated as a speculative demonstration of the underlying concept, with limitations that are left for future work.

In Figure Set 3 below (medium grey bars) we suggest that, relative to unselected cases, using severity enriched cases can inflate estimates of liability scale heritability, relative to unascertained cases. As above, we similarly use equation 4 to back calculate the necessary K_U_ to return the heritability estimate provided in the unascertained scenario, providing a benchmark estimate for bounding potential impact of sampling extreme cases. In Figure Set 3 (dark grey bars) we show how using both extreme cases and controls compounds the effect introduced by each individual factor, as expected. Intuitively, we can interpret the estimates of K_L_ and K_U_ as demonstrating that extreme sampling of cases and controls can lead to a “gap” in a study’s representative of the underlying population liability, which should follow the implications of Yap et al.

In Figure Set 4 below we show the corrected external study SNP heritability estimates that attempt to account for various levels of extreme sampling of controls or cases, independently, adapting the framework of Yap et al^21^. These corrections using the bounds for K_L_ and K_U_ we derived from resampling iPSYCH cases or controls suggest the heritability estimates could potentially be fairly substantially biased upwards (black dot and confidence interval vs. the grey dot and confidence intervals, along the continuous colored lines). In Figure Set 5 below we show corrections for nine permutations assuming *both* extreme cases (K_U_ = K*p, p=0.1, 0.5, 0.75) *and* extreme controls (K_L_ = (1-K)*p, p=0.75, 0.85, 0.95), where our scaling factors, p, were chosen to span the plausible ranges gleaned from Table 4, below. Even positing among the weakest scenarios is enough to remove the differences between external estimates and those made in iPSYCH for some disorders (ADHD, AFF, ANO, XDX), although quite strong selection in both directions is needed for others (ASD, BIP, SCZ).

**Figure Set 5.**
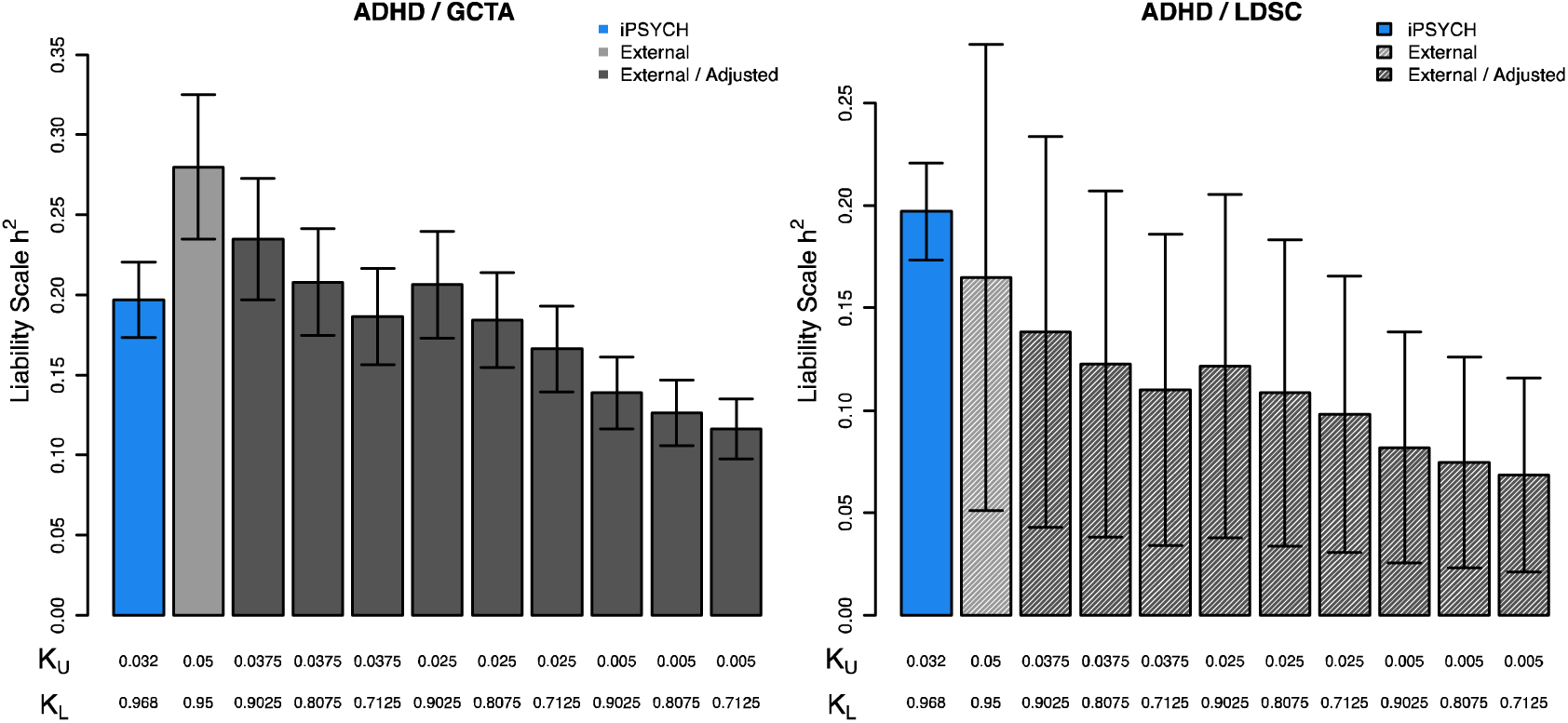
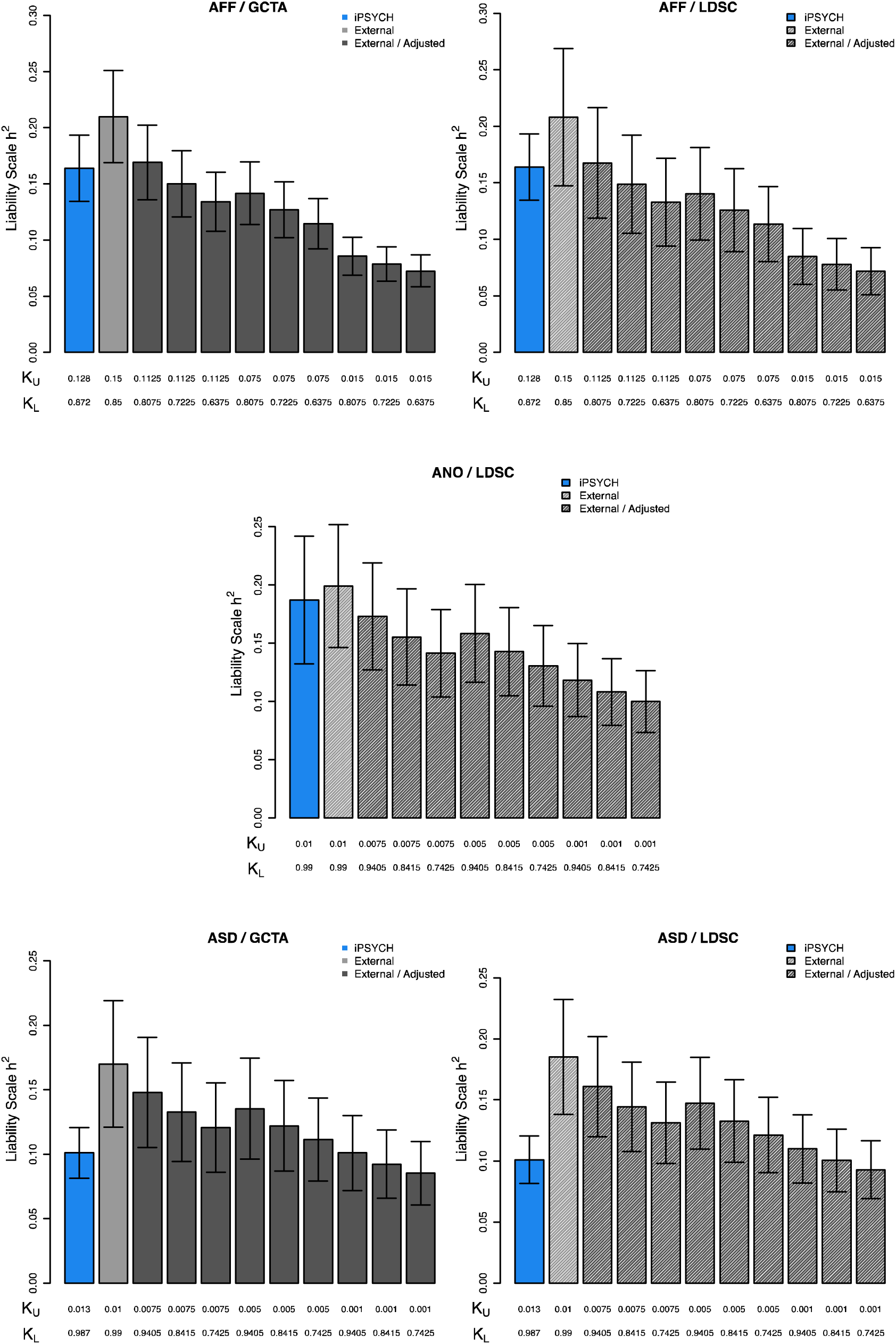
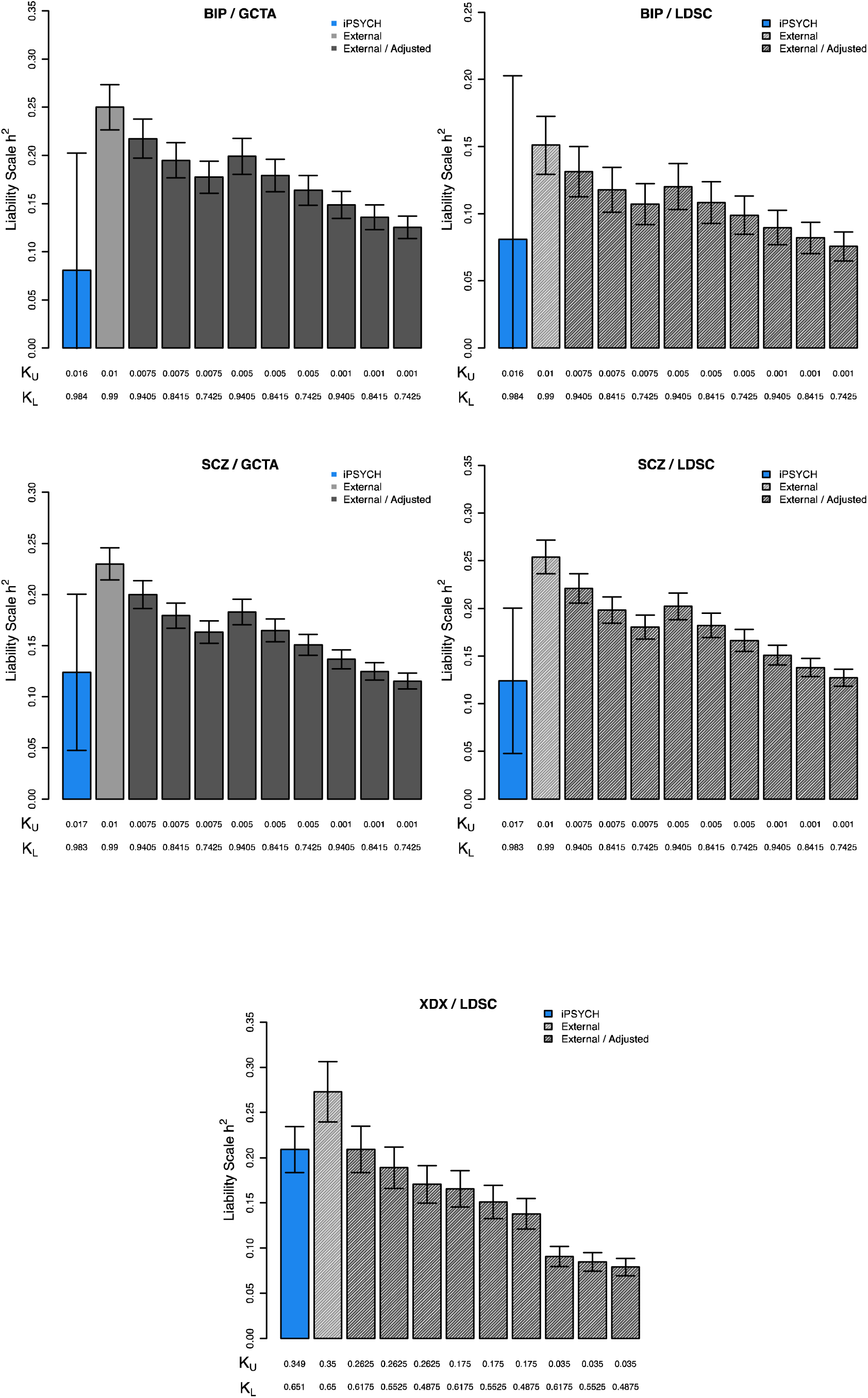
Exploring the potential effect of simultaneous extreme sampling of cases and controls for a limited number of scenarios. Each pair of figures (row) presents the unadjusted iPSYCH estimate of SNP heritability (blue) and external study (light grey) along with estimates corrected for one of nine permutations of K_U_ = K*p where p=0.75, 0.5, or 0.1 and K_L_ = (1-K)*p where p=0.95, 0.85, 0.75 which more or less cover the values from Table 2 above. GCTA estimates are shown with solid bars and LDSC estimates with shaded bars.

As was described by Yap et al, when cases and controls have been selected to occupy more extreme portions of the underlying population liability, substantial bias in heritability can arise, however, precise estimates of the size of the effect require precise study into the strength of extreme sampling. We emphasize that this exercise and our attempts to bound potential overestimation from external estimates of SNP-heritability for psychiatric conditions in data that has been ascertained for GWAS are limited and should be seen as speculative. However, we do feel it suggests at least *some* overestimation of heritability should be expected in the published studies, given our knowledge of their ascertainment schemes. More precise investigations into the extent of overestimations, and whether the effects have a substantial impact on our broader conceptualizations of the genetic architecture of psychiatric conditions, is a topic for future research, as is the effect of extreme sampling on SNP-based genetic correlations, for which we are unaware of an established framework for investigating.

### Diagnostic Errors

Our interpretation of statistics associated with a disease (observed disease) may differ from our belief of the disease (true disease). When cases are ascertained through noisy diagnoses, the estimated of SNP-heritability from the observed disease are typically downward biased compared to the *true* disease^26^, of course ascertaining on true disease status may never be achievable. For example, if some proportion of cases of one disorder are instead diagnosed with a different disorder, and/or vice versa, then estimates of genetic correlations between these two disorders will be inflated^26^. Wray et al.^26^ introduced a framework for exploring the expected impact of various levels of diagnostic error for two hypothetical disorders, A and B, in terms of unstandardized additive genetic variances and covariance for the diagnosed 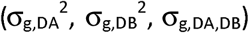 and true disorders 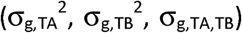, and the proportions of cases correctly diagnosed (M_TA_, M_TB_) and misdiagnosed (M_FA_=1-M_TA_, M_FB_1-M_TB_). Antilla et al^27^ re-derived the single variance component 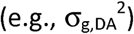 equation in terms of liability scale heritability (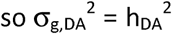 as the liability variance is taken to be one) and defining the covariance term (σ_TA,TB_) using the genetic correlation 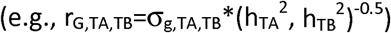, giving an equation for describing the relationship between the true heritability and that estimated from diagnosed individuals.

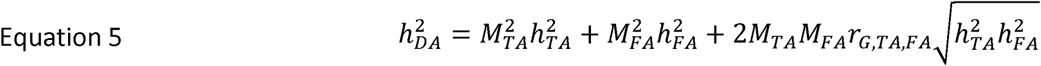

Under the simplifying assumption that for a given pair of disorders, the misclassification occurs only between the two disorders for which the correlation is computed, such that the misdiagnosed proportion of those diagnosed with disorder A (M_FA_) in actuality have disorder B, we can use the same notation to re-write the equations for σ_g,DA,DB_ from Wray et al^26^ in terms of genetic correlations and liability scale heritability.

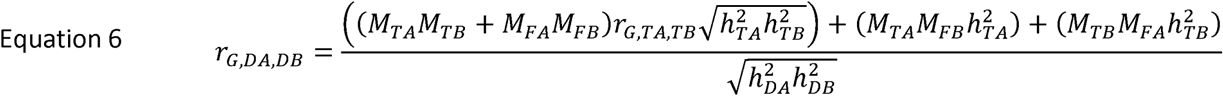

From our knowledge of the external meta-studies, the contributing cohorts are predominantly ascertained according to research diagnoses, while in in iPSYCH we aggregate register records of clinical diagnoses. Although psychiatric diagnoses in the Danish register have been shown to be highly reliable in multiple validation studies (e.g.,^28-31^) the diagnoses in iPSYCH cohort, *per se*, have not been systematically re-assessed and the trends towards lower SNP-heritability and higher SNP-based genetic correlations could, in theory, be taken as evidence for a relative increase in misdiagnosis when compared to the rate in external studies. Previous work in the Danish health registers^32^ has suggested that ~15% of patients first diagnosed with BIP will be subsequently diagnosed with SCZ and ~6% initially diagnosed with SCZ will be subsequently diagnosed with BIP, numbers that more or less agree with an independent study^33^ conducted in the U.S. (~15% and ~4%). We can speculate whether these shifts represent initial misdiagnoses, disease progression, comorbidity, or some other factor, and the extent to which they also affect external meta-studies (an un-measureable phenomenon, given the lack of lifetime data). Regardless, we can anchor our investigations into the magnitude of effects of potential misdiagnosis on a range with some empirical support (~5-15%), which may represent an upper bound given the similarities between the symptoms of SCZ and BIP, relative to other disorders, and because we attribute the entire cause of these later diagnoses to initial misdiagnoses.

In Figure set 6, below, we investigate the potential effects of various levels misdiagnosis on estimates of SNP-heritability. In order to predict a “true” SNP-heritability 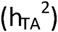 from a “diagnosed” SNP-heritability 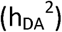, we need to specify not only the misdiagnosis rate (M_FA_), but also the composite SNP-heritability of the disorders the misdiagnosed patients truly have (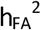, 0 if they should be a control), and the genetic correlation between the true patients and misdiagnosed patients (r_G,TA,FA_). We consider three values for each which span the range of estimates in Table 1 (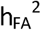 = 0, 0.1, 0.2; r_G,TA,FA_ = 0, 0.33, 0.66). Under the intuition that a patient diagnosed is unlikely to be a true control but may have been assigned the wrong disorder, we place special emphasis on the dashed blue and purple lines, which consider the misdiagnosed patients to be a group that has moderate genetic correlation with the disorder of interest, consistent with an “average psychiatric patient.” These data suggest that misdiagnosis rates ranging from 5-15% can lead to a modest under estimation of SNP-heritability, but unreasonably high misdiagnosis rates (>40%) are needed to rectify the largest differences. For some disorders and parameter scenarios (e.g., BIP, purple lines), misdiagnosis alone cannot, even in theory, explain the observed differences in SNP-heritability, no matter the proposed rate.

**Figure Set 6.**
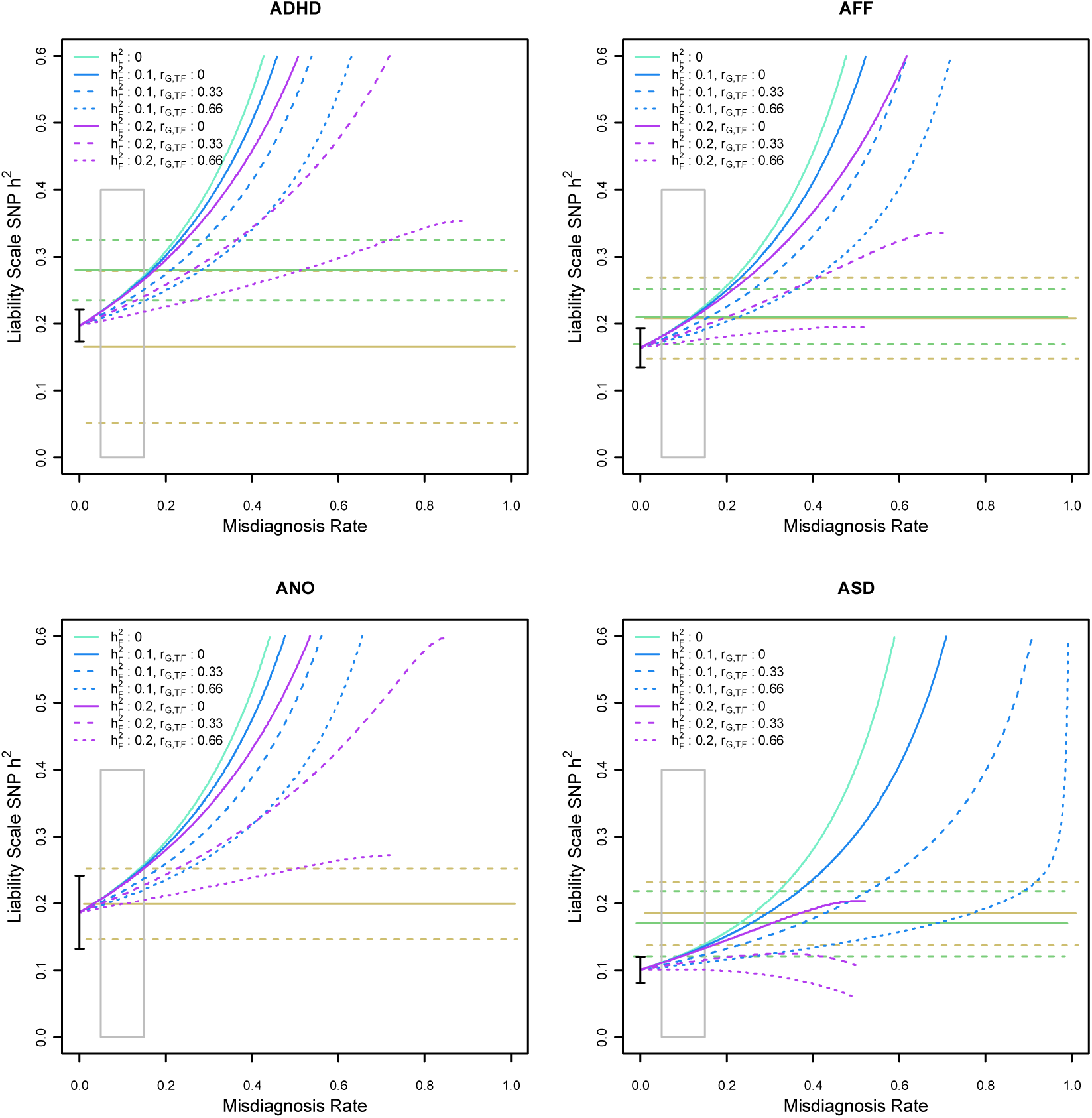
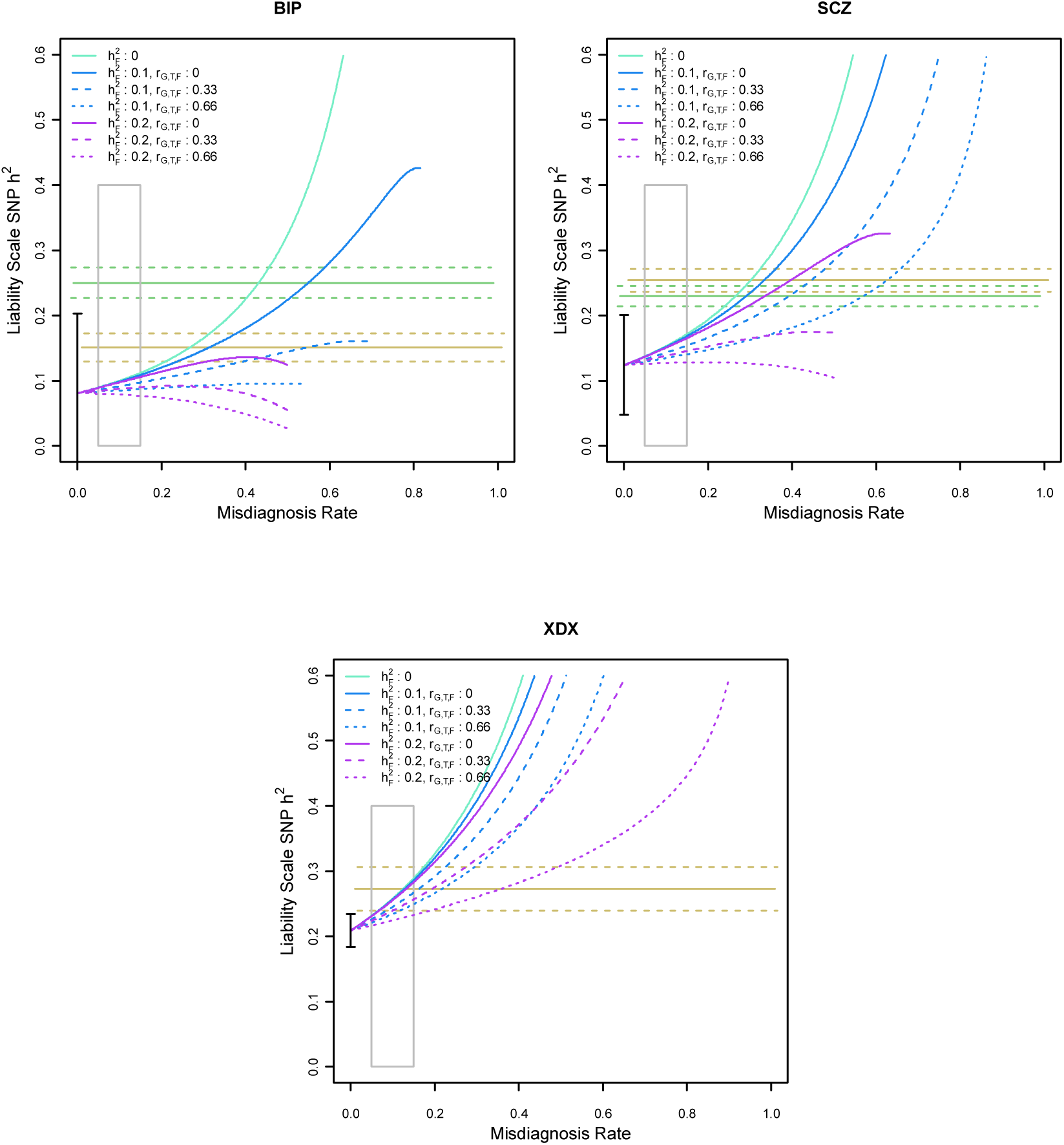
For each iPSYCH disorder we attempt to describe the potential contributions of misdiagnosis. Values on the y-axis represent the “true SNP heritability” that would be expected produce our estimate from Table 1, under varying levels of misdiagnosis (x-axis). We consider scenarios where the misdiagnosed patients represent a group with a SNP-heritability of 0 (aqua lines), 0.1 (blue lines), or 0.2 (purple lines) which is genetically correlated either 0 (solid lines), 0.33 (dashed lines) or 0.66 (dotted lines) with the diagnosis of interest. Yellow and green rectangles show the 95% confidence intervals for the LDSC regression and GCTA external estimates of SNP-heritability, respectively, presented in Table 1. The grey box highlights a 5-15% window. Black dot and bars represent point estimate and 95% confidence interval from Table 1.

Unlike the previous examples, the effects of misdiagnosis on SNP-based genetic correlations *has* been described analytically, albeit only for the scenario where cases from the two disorders for which genetic correlations are being estimated are misdiagnosed for each other, only. In Figure set 7, below, we show the levels of inflation that are expected for a given true value of genetic correlation (green bars), when various proportions of the two hypothetical disorders are misclassified as the other (grey bars). When true genetic correlations are low (0 or 0.2) and misdiagnosis rates are high (0.15), the genetic correlations can be exaggerated by nearly 0.40 above their true value. When true genetic correlations are more moderate and misdiagnosis rates lower (0.05), the expected inflation is more modest.

**Figure Set 7.**
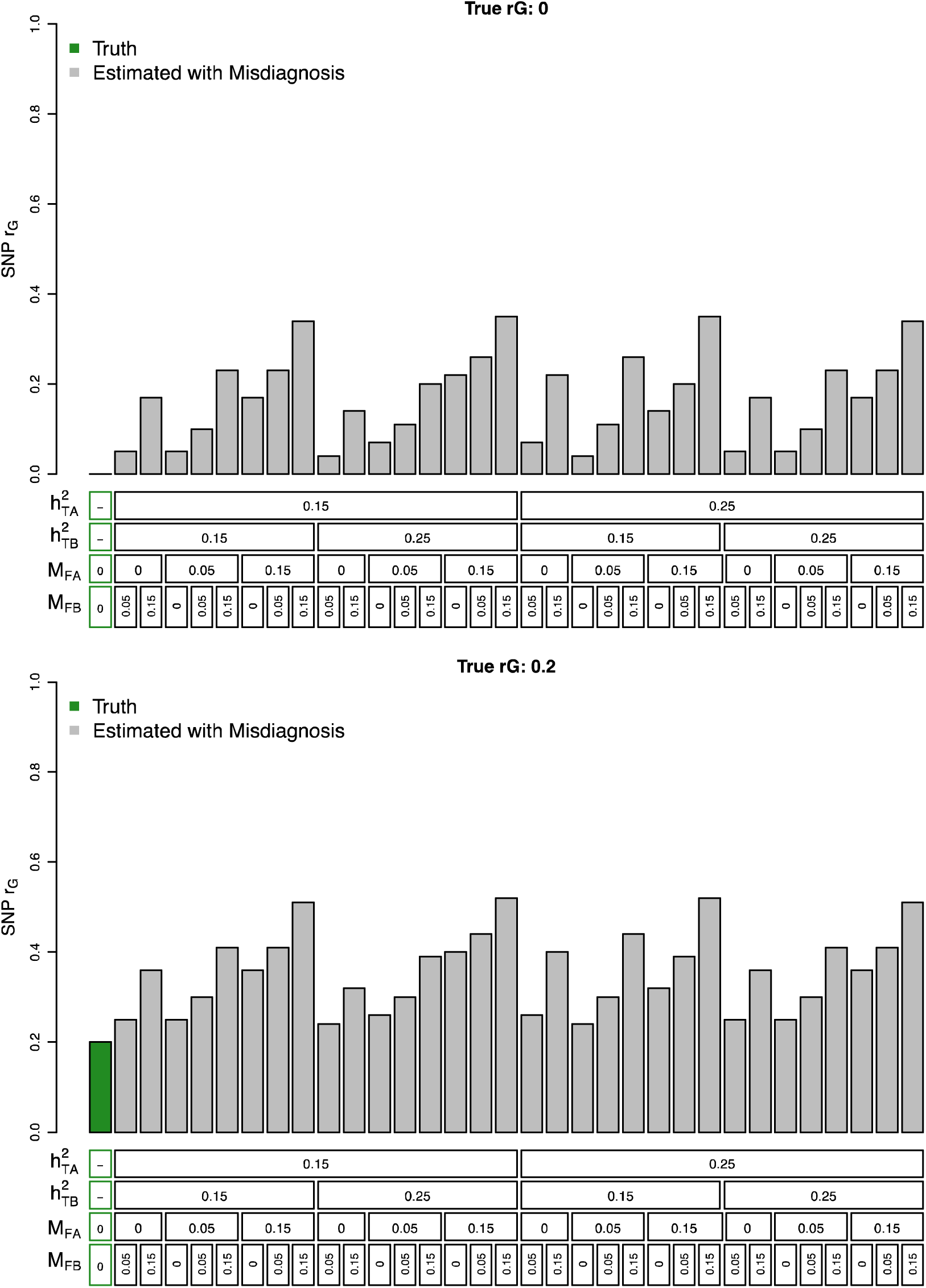
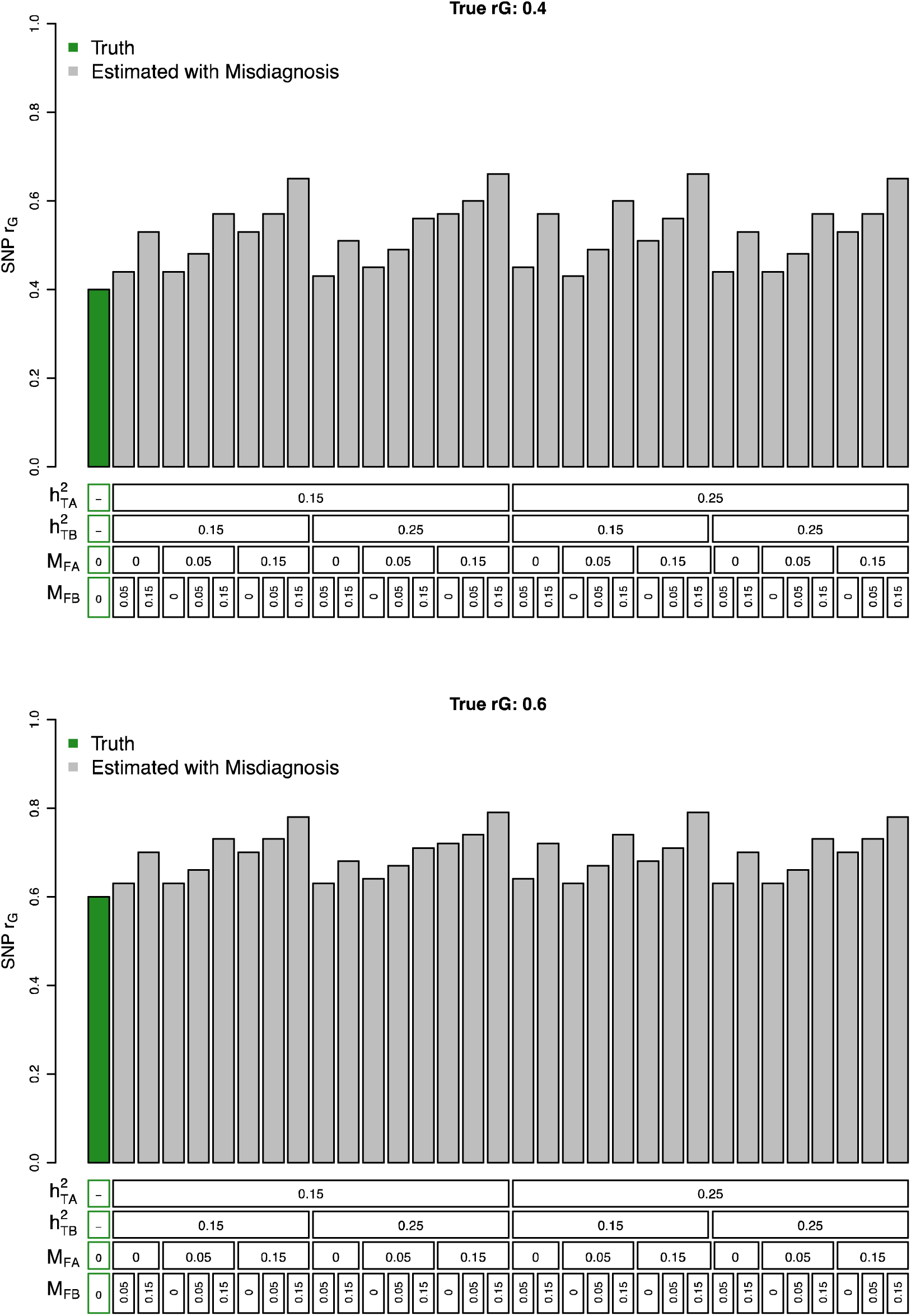
Potential impact of diagnostic misclassification on estimates of SNP-based genetic correlations. For four levels of “true” genetic correlation (True r_G_ = 0, 0.2, 0.4, 0.6) and two levels of true heritability for each hypothetical disorder 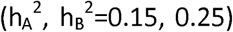, we describe the predicted effects of different combinations of misdiagnosis 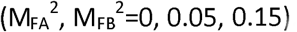. The green bar shows the “true” genetic correlation, while the grey bars show this inflation due to different combinations of true SNP-heritability and levels of misdiagnosis. Note that all misdiagnosed cases of A are assumed to be true cases of B, and vice versa.

## Discussion

In this work we have, where established frameworks were available, attempted to provide context for the potential sources of uncertainty in SNP-based heritability and genetic correlation estimates that could contribute to subtle difference trends between iPSYCH and external meta-study reports. In doing so, we hope to emphasize a few points. First, strong interpretations of exact point estimates for SNP-heritability and genetic correlations may impose more precision than the underlying models and concepts beget. We see these quantities as providing broad-scale benchmarks for more qualitative claims regarding the amount of genetic variation remaining to be discovered by GWAS, general importance of common SNP variation and the plausibility of pleiotropic signals among risk variants. Second, ascertainment of cases and controls may be an important consideration when estimating SNP-heritability and one that could be studied further. Here, we consider two sets of studies, iPSYCH and external meta-studies, which were ascertained according to different sampling schemas (registers vs. predominantly clinical ascertainment) and with different primary goals in mind (epidemiological validity vs. GWAS power). As we have attempted to demonstrate, the differences in these schemas may make the individuals studies susceptible to different sources of bias that may not be a part of the current intuition when considering estimates of SNP-heritability and genetic correlations. More work in this area may be needed as large genetic cohorts with different ascertainment schema emerge.

We can summarize the totally of these demonstrations by integrating them with our own intuition about the most likely consequences of differences in ascertainment between iPSYCH and external studies. Taken together, we could speculate that the variability in SNP-heritability estimates from the meta studies is likely underestimated due to variability in the assumed lifetime risk that is not accounted for, may be underestimated modestly in iPSYCH due to the relatively younger age of control subjects and potentially higher incidence of misdiagnosis, and overestimated in external studies due to the use of extreme cases and controls. In terms of genetic correlations, we are unaware of frameworks for exploring the potential effects of age or severity censoring, but increased misdiagnosis rates could lead to upwardly biased estimates of genetic correlations. These conclusions should be viewed against the background of the inherent lack of precision in the models and concepts of SNP-heritability and genetic correlations, large sampling variances for the estimators, expectations for real differences when genetic parameters are estimated in different populations, and the need for future development of frameworks for further studying the effects of ascertainment on genetic parameters.

